# The DNA methylation enzymatic machinery in substance use disorders: a systematic review

**DOI:** 10.1101/2025.11.12.688056

**Authors:** Margot Diringer, Mathieu Bruggeman, Pierre-Eric Lutz

## Abstract

Substance use disorders (SUD) are chronic affections defined by similar symptoms across a variety of psychoactive drugs, including alcohol, cocaine, opioids, or methamphetamine. Epigenetic mechanisms such as DNA methylation represent key candidates to help explain the long-lasting effect of these drugs, as well as inter-individual variation in vulnerability. Here, we systematically reviewed current knowledge on the role of DNA methylation and the related enzymatic machinery in rodent models of SUD. Using a prospectively registered methodology, 99 articles were prioritized. A first set of studies manipulated the expression or activity of methylation or demethylation pathways. Depending on the brain region or drug considered, SUD-related behavioral and molecular manifestations were bidirectionally modulated, suggesting both pathogenic and protective roles for drug-induced methylomic plasticity. A second set of articles focused on candidate genes. Although significant heterogeneity across experimental models, brain regions or gene targets resulted in an absence of replicated findings, available data nevertheless support the notion that drugs of abuse trigger DNA methylation changes at discrete loci. Third, recent genome-wide studies have started to demonstrate that these drugs recruit widespread reprogramming. Strikingly, most adaptations occur outside promoter regions, highlighting an important challenge toward their functional interpretation. Finally, studies of drug exposure during gestation or adolescence suggest long-lasting consequences, with the potential for early intervention.

## 1. Introduction

Substance use disorders (SUD) are chronic affections defined by similar symptoms across a variety of psychoactive drugs, such as cocaine, opioids, alcohol, or THC (Hasin et al., 2013). These symptoms include excessive motivation to get the drug, persistent use despite harmful consequences, or uncontrollable drug seeking. Similar features are shared with behavioral addictions, defined by uncontrolled repetition of reinforcing behaviors (Robbins and Clark, 2015). Despite intensive research, the neurobiological bases underlying the emergence or the protracted course of these behaviors remain poorly defined. Notably, we still ignore why only a minority of individuals who use drugs for recreational or therapeutic purposes develop an addiction, or why those who manage to achieve abstinence face a lifelong risk of relapse.

Recently, epigenetic mechanisms have been proposed to contribute to such inter-individual variability or prolonged clinical course (Falconnier et al., 2023; Hamilton and Nestler, 2019). Epigenetics, which can be defined as the physical and chemical processes that control the organization and functional expression of the genome, notably include histone modifications, non-coding RNAs and DNA methylation (DNAm). DNAm, first reported in bacteria a century ago (Johnson and Coghill, 1925), is involved in a wide variety of biological processes. These include genomic stability, developmental regulation of gene expression, transposon silencing, imprinting, or X chromosome inactivation (Greenberg and Bourc’his, 2019; Schübeler, 2015). Such processes are mediated by dynamic changes in the abundance and location of DNAm along the genome, which, in turn, are dependent on enzymatic pathways responsible for methylating and demethylating the DNA (Dnmt and Tet, respectively, see below). More recently, these pathways have started to be implicated in neuropsychiatric diseases, including SUD. In this context, the goal of the present systematic review was to analyze current knowledge on the role of DNAm and the related enzymatic machinery in SUD.

To do so, we used a preregistered methodology to identify studies that investigated the impact of manipulating these enzymes on behavioral and molecular effects of drugs of abuse, or characterized genomic patterns of DNAm following exposure to these drugs. Below, we first provide a brief overview of current knowledge on biological functions and mechanisms of action of DNAm in brain tissue. Second, based on our bibliographic screening of rodent studies, we provide a qualitative synthesis of available data, and review experimental designs and main findings. Finally, we discuss perspectives for future investigation in the field.

## 2. The brain DNA methylation enzymatic machinery

DNAm exists in the form of 5-methylcytosine (5mC), N6-methyladenine (6mA) and N4-methylcytosine (4mC). As 4mC has not been described in eukaryotes, and the existence of 6mA remains controversial (Alderman and Xiao, 2019), we focused in the present review on 5mC. While this type of DNAm is mostly found at CG dinucleotides in vertebrates (Sinsheimer et al., 1954), early work in other organisms has shown that it also affects additional contexts, such as CHG and CHH in plants (where H stands for A, C or T (Pikaard and Mittelsten Scheid, 2014)). Similar non-CG DNAm (5mCH) has more recently been characterized in mammals (He and Ecker, 2015), where it is mostly restricted to a few cell types that include embryonic stem cells or myocytes, as well as neurons, in which they reach unexpectedly high levels (see (de Mendoza et al., 2021) and below).

Genomic patterns of DNAm result from combined activities of methylation and demethylation pathways that implicate a limited number of enzymes, contrasting with the numerous actors involved in histone post-translational modifications. These patterns are established during embryonic development and cellular differentiation by 2 DNA methyltransferases, Dnmt3a and Dnmt3b, that perform *de novo* methylation (Okano et al., 1999). During mitotic divisions, maintenance of DNAm is ensured by Dnmt1, which shows high affinity for hemimethylated CG sites generated during DNA replication (Fatemi et al., 2001; Hermann et al., 2004). The second component of the DNAm cycle is its removal. Passive demethylation occurs when newly synthesized DNA strands go unmethylated over successive cell divisions, while active demethylation is catalyzed by 3 Ten Eleven Translocation enzymes (Tet1-3). These enzymes successively oxidize 5mC to 5-hydroxymethylcytosine (5hmC), 5-formylcytosine (5fC) and 5-carboxylcytosine (5caC) (Tahiliani et al., 2009; He et al., 2011; Ito et al., 2011), ultimately leading to their replacement by unmodified cytosines via base excision repair. Among these demethylation intermediates, only 5hmC has been found stable in specific cell types, reaching its highest levels in the central nervous system (Kriaucionis and Heintz, 2009), with about 8, 12 and 20% of CG sites hydromethylated in microglia, astrocytes and neurons, respectively (Tooley et al., 2023; Wei et al., 2024).

DNAm landscapes vary significantly across tissues and cell-types. Genomic regions with accessible chromatin and high transcriptional activity tend to be lowly methylated, coherent with the notion that DNAm opposes the binding of transcription factors (Stadler et al., 2011), in particular at promoters and enhancers (Luo et al., 2018). More than two-thirds of mammalian promoters also harbor CG islands (CGI), which correspond to ∼1 kb-regions with high CG density, where bimodal methylation levels (low/high) act as on/off signals for transcription (Bird et al., 1985). In contrast, DNAm levels are typically higher and less variable in other genomic regions such as satellite DNA, repeated elements, or transposons (Li and Zhang, 2014). Downstream functional consequences of DNAm are primarily mediated by the recruitment of proteins that bind methylated CG sites (via their methyl-CpG binding domain, MBD), as well as interactions with other epigenetic mechanisms (e.g. histone modifications). Proteins of the MBD family include MeCP2, which is mutated in the neurodevelopmental Rett syndrome (Liu et al., 2025; Tillotson and Bird, 2020), as well as Mbd1-4 (Bird and Wolffe, 1999). These proteins in turn interact with histone modifying enzymes, such as deacetylases (HDACs), thereby modulating other epigenetic layers to regulate transcription.

Over the last decade, several specificities of the brain epigenome have been uncovered. Among these is the fact that neurons are post-mitotic cells that exhibit a progressive and strong accumulation of 5mCH and 5hmC, at much higher levels than glial cells, with potentially significant implications for psychopathology (Lutz et al., 2021, 2017). The accumulation of 5hmC starts in utero (Lister et al., 2013) and continues after birth, throughout aging (Szulwach et al., 2011). For 5mCH, the process starts later and primarily occurs during the first few weeks (mouse) or years (human) of postnatal life (Lister et al., 2013). Interestingly, the neuronal pattern of 5mCH has been involved in the repression of long (>100 kb) genes through the recruitment of MeCP2 (Gabel et al., 2015; Moore et al., 2025; Tillotson et al., 2021). This postnatal accumulation is mediated by enhanced expression and activity of Dnmt3a (Stroud et al., 2020, 2017), and mostly affects CAC sites, in contrast with embryonic stem cells, in which non-CG methylation occurs at CAG (He and Ecker, 2015). This central role of Dnmt3a in shaping the brain methylome reflects the fact that, although Dnmt3a and Dnmt3b cooperate to mediate waves of DNAm during embryological development, the latter is lowly expressed in the brain, while expression of the former peaks during the postnatal period and remains high thereafter (Lister et al., 2013). While we now have a detailed understanding of these kinetics, the persistence in the adult brain of an active turnover involving both methylation and demethylation of the DNA remains an open question.

## 3. Methods of the systematic review

### 3.1 Identification of studies

The review protocol was registered prospectively in Prospero (CRD42024440109; https://www.crd.york.ac.uk/prospero/display_record.php?RecordID=440109), and followed the Preferred Reporting Items for Systematic Review and Meta-Analysis protocol, PRISMA (Liberati et al., 2009). A systematic electronic search was conducted in 3 databases: PubMed, Web of Science and Scopus (Fig.1). This search was performed until August 2023, with no restriction on publication date, using the following keywords: “Dnmt” OR “DNA methyltransferase” OR “DNA methylation” OR “Tet” OR “ten eleven translocation” AND “addiction”. The titles of all articles were screened to remove duplicates.

**Figure 1.**
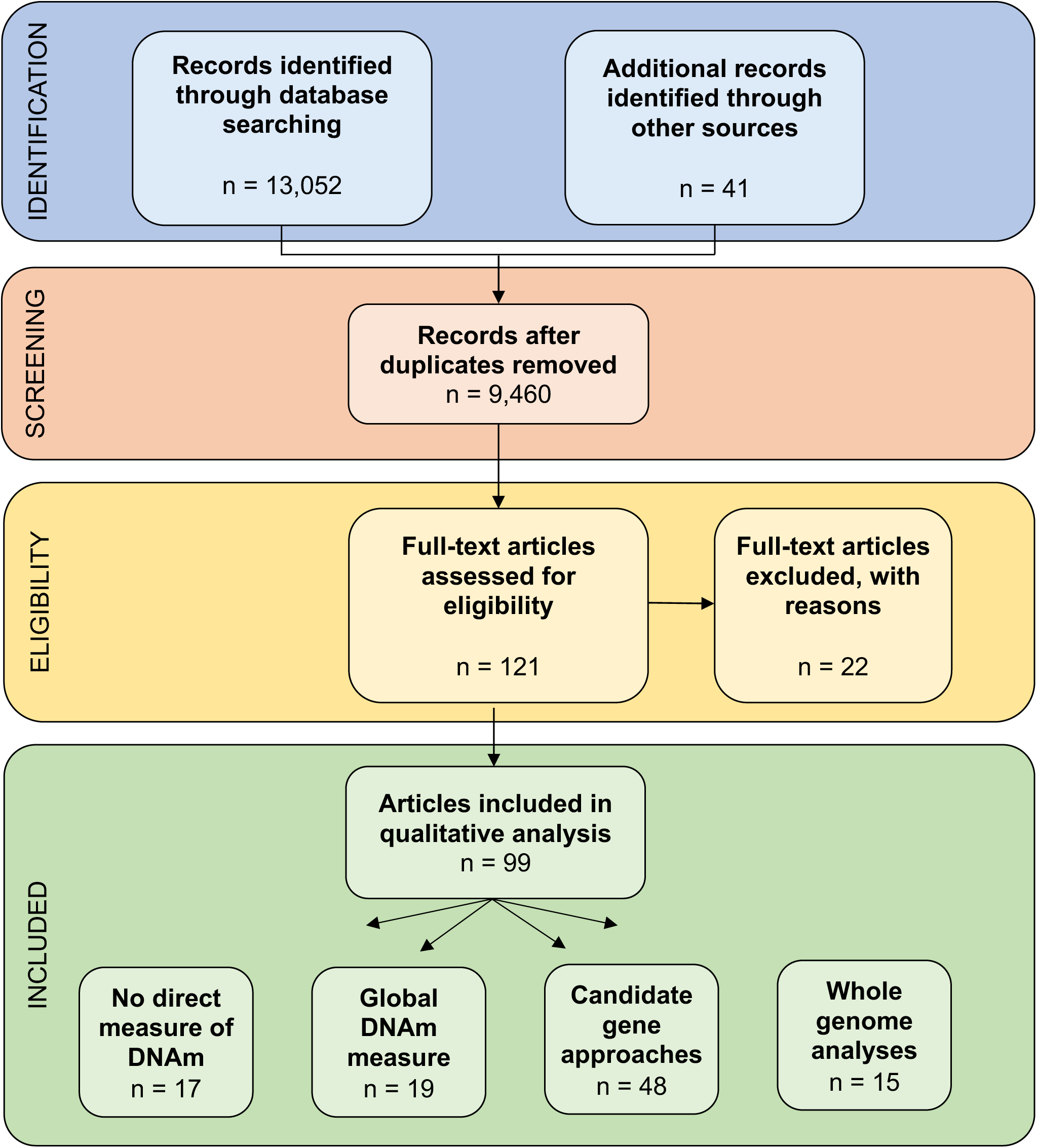
PRISMA diagram. The review protocol was registered prospectively in Prospero (record ID# 440109), and followed the Preferred Reporting Items for Systematic Review and Meta-Analysis protocol (PRISMA (Liberati et al.,2009)). *Identification of studies*. Three databases were used for systematic screening: PubMed, Web of Science and Scopus. The literature search was performed using the following keywords: “Dnmt” OR “DNA methyltransferase” OR “DNA methylation” OR “Tet” OR “ten eleven translocation” AND “addiction”, providing 13,052 potentially eligible articles. Forty additional records identified through other sources were also considered. *Screening*. Articles were screened based on their title and abstract, and included if they interrogated the link between enzymes responsible for DNAm or demethylation and molecular or behavioral effects of drugs of abuse (opioids, cocaine, alcohol, THC, nicotine or amphetamine; see *Methods*). Duplicates were also removed. *Eligibility*. After a more thorough examination of the full-text of the 121 remaining articles, an additional 22 articles were excluded. As a result, 99 papers were selected for qualitative synthesis and organized according to the experimental approach (see main text for details).

**Figure 2.**
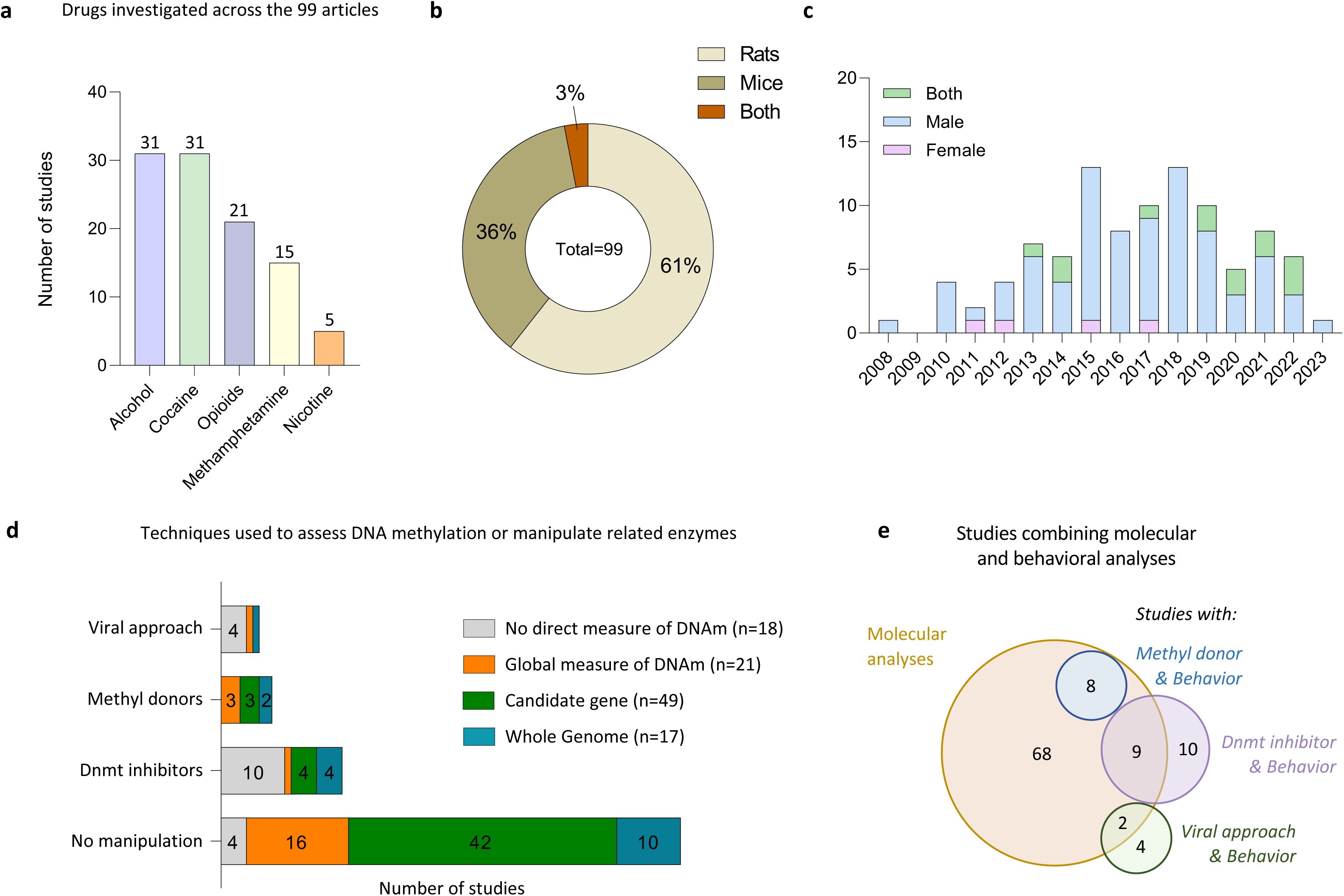
Characteristics of studies included. **a.** Drug of abuse investigated in the 99 included articles (NB: 4 articles studied 2 drugs (Tian et al., 2012; Fragou et al., 2013; Chao et al., 2014; Mychasiuk et al., 2013). **b.** Distribution of rodent species across studies. **c.** Distribution of sex and year of publication across studies. **d.** Techniques used to assess DNA methylation or manipulate related enzymes (NB: 4 articles used 2 types of manipulations (Qiang et al. 2014; Massart et al. 2015; Zhang et al. 2020; Hong et al. 2020), and 1 used 3 (LaPlant et al., 2010). **e.** Venn diagram of the number of studies that combined molecular and behavioural analyses and used different strategies to manipulate DNA methylation enzymes (Methyl donor, Dnmt inhibitor, viral approach).

### 3.2 Eligibility

Articles were assessed for eligibility based on their title and abstract. When not sufficient, the main text was also screened. Studies were considered eligible if they interrogated the impact of drugs of abuse on DNAm (opioids, cocaine, alcohol, THC, nicotine or amphetamine), or the link between enzymes responsible for DNAm and molecular or behavioral effects of these drugs. The literature search was restricted to rodent studies, published in English, in peer-reviewed journals. Exclusion criteria were: (i) studies conducted in humans, *in vitro*, using cell lines, or in tissues other than the nervous system; (ii) the absence of a control group ; (iii) or a focus on diseases associated with drug exposure other than SUD (e.g cancer).

## 4. Description of eligible articles

A total of 99 articles met our inclusion criteria, among which 5 drugs of abuse were represented, with alcohol (n=31, 31%) and cocaine (n=31, 31%) the 2 substances most frequently investigated (Fig.2a). Four articles studied 2 substances, corresponding to a total of 103 studies. Most of these were conducted in rats (61%) and in mice (36%), and few investigated both species (3%) (Fig.2b).

Regarding sex, studies in male rodents were largely predominant (n=81, or 83%, as opposed to 17% for those conducted in females; Fig.2c), although a trend towards a consideration of both sexes emerged over the last few years. These numbers are similar to those observed in previous reviews in molecular psychiatry (Falconnier et al., 2023; Beery et al., 2011). Importantly, while epidemiological data indicate that SUD are more frequent in men, they nevertheless also represent a public health issue in women, who develop the condition under the influence of partly distinct risk factors (Becker and Chartoff, 2018) and exhibit different symptomatic profiles (Nicolas et al., 2022). As such, there is an urgent need to move toward a more systematic analysis of sex differences.

Two types of studies were distinguished based on whether they conducted molecular measures of DNAm or manipulated the DNAm enzymatic machinery. Among the 99 articles included, some used more than one approach, corresponding to a total of 104 experiments. A majority of these studies (n=86/104; 83%) conducted DNAm measures either: (i) globally, at genome-wide level, using e.g. ELISA approaches (n=21, 24%; Fig.2d, orange cells); (ii) for specific candidate genes (n=49, 57%; green cells); or (iii) using methods that assessed cytosines throughout the genome (by microarrays or sequencing; n=17, 20%). Surprisingly, only 19 experiments combined DNAm measures with a manipulation of the machinery (18%; Fig.2e), indicating that the 2 strategies are rarely used together. In contrast, the experiments that involved manipulating the enzymes represented a minority (n=33/104; 32%), and used either: (i) pharmacological inhibitors (n=19); (ii) viral strategies to modify their expression (n=6); (iii) or methyl donors to increase DNAm levels (n=8). Finally, a small corpus of articles (n=4) used neither approaches, and instead quantified the expression of DNAm enzymes as a function of drug exposure.

## 5. Manipulation of the activity of DNA methylation enzymes

Three pharmacological agents have been used to inhibit Dnmt enzymes: 5-aza-2ʹ-deoxycytidine (5-aza), zebularine and RG108. Initially developed for cancer treatment, 5-aza is incorporated into DNA during replication and forms covalent bonds with the enzymes. This triggers their degradation, leading to progressive DNA hypomethylation and reactivation of tumor suppressor genes (Stresemann and Lyko, 2008). Zebularine is another analog of cytidine with a similar mode of action. While these compounds have proved potent in dividing cancer cells, their impact on post-mitotic neurons remains comparatively less well characterized, but may involve neurotoxicity (Wang et al., 2013). On the other hand, RG108 was the first synthetic compound designed to reversibly inhibit the catalytic domain of Dnmt enzymes, leading to DNA demethylation and reactivation of tumor suppressor genes, similar to 5-aza. In the brain, a single study showed that RG108 reduced DNAm, an effect supported by an ELISA-like measure of global genome-wide levels (LaPlant et al.,2010). In addition to Dnmt inhibition, a second approach is to administer methyl donors, typically via dietary supplementation (either S-adenosyl-L-methionine, SAM, or its precursor methionine). Because such donors affect all methylation reactions within cells (Gao et al., 2018), their impact on the transcriptome and methylome are likely complex, bidirectional, and may affect multiple loci (Wang et al., 2017). In the context of SUD, these effects remain to be more thoroughly characterized, as detailed below. Third, viral strategies have been used to overexpress or knockdown Dnmt or Tet enzymes. Compared to pharmacological or dietary approaches, these strategies enable targeting more specifically a single enzyme or cell-type of interest, although they are more difficult to implement. In the following sections, we summarize how they have been used to study ethanol, cocaine, and opioids. Experimental designs and behavioral results are summarized for each drug in Fig.3, while Fig.4 differentiates studies that argue for a pathogenic or protective role of Dnmt and Tet enzymes in SUD-related phenotypes.

**Figure 3.**
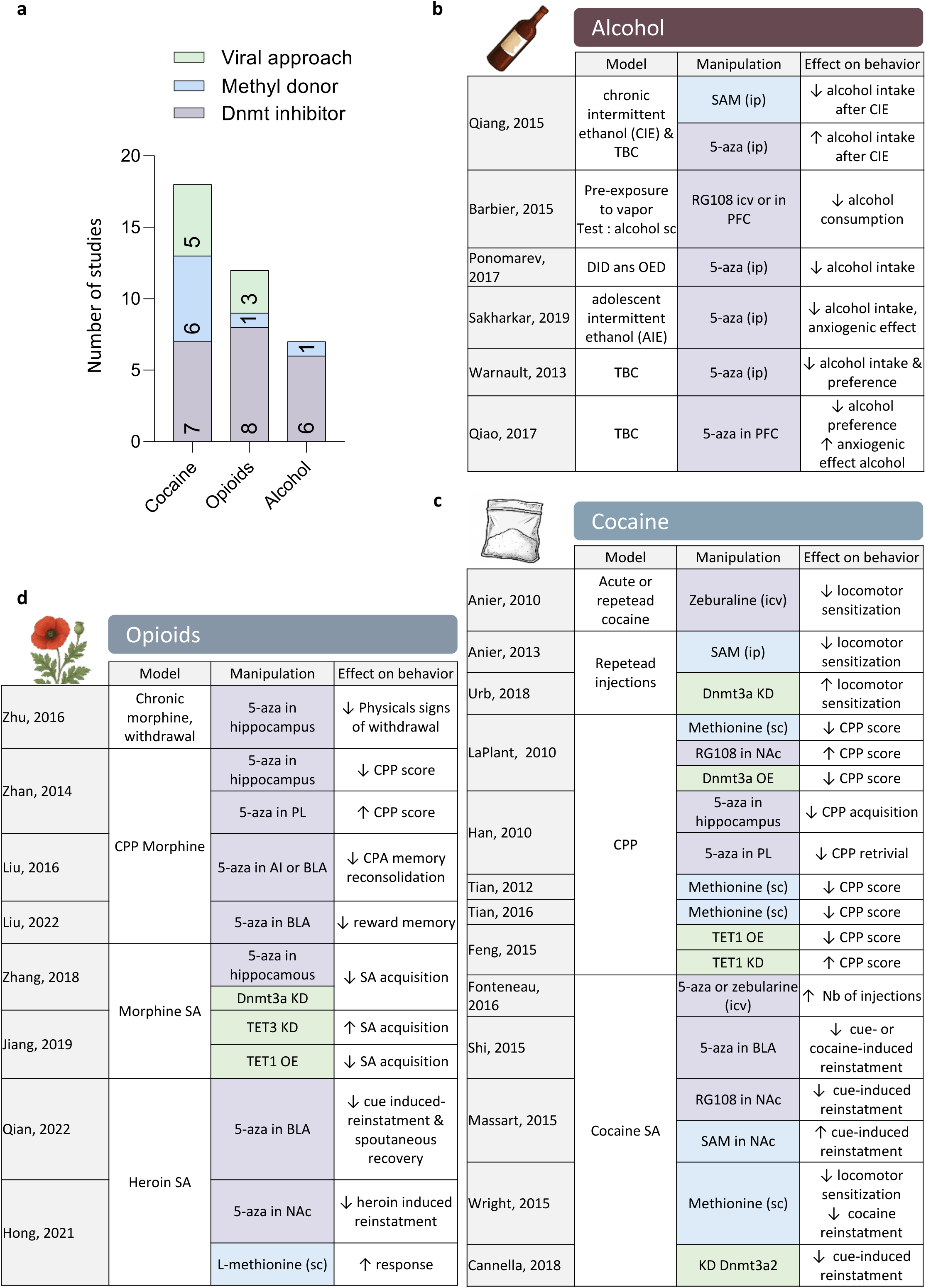
Summary of experimental methods and results from studies that manipulated DNA methylation enzymes. **a.** Number of articles according to the drug of abuse under study and type of manipulation. **b-d.** Summaries of behavioral models, type of manipulation and results obtained for studies that investigated alcohol (b), cocaine (c) or opioids (d). Colours indicate the method used (green, viral approach; blue, methyl donor; purple, Dnmt inhibitor). Upward and downward arrows indicate an increase or a decrease in behavioral responses, respectively.

**Figure 4.**
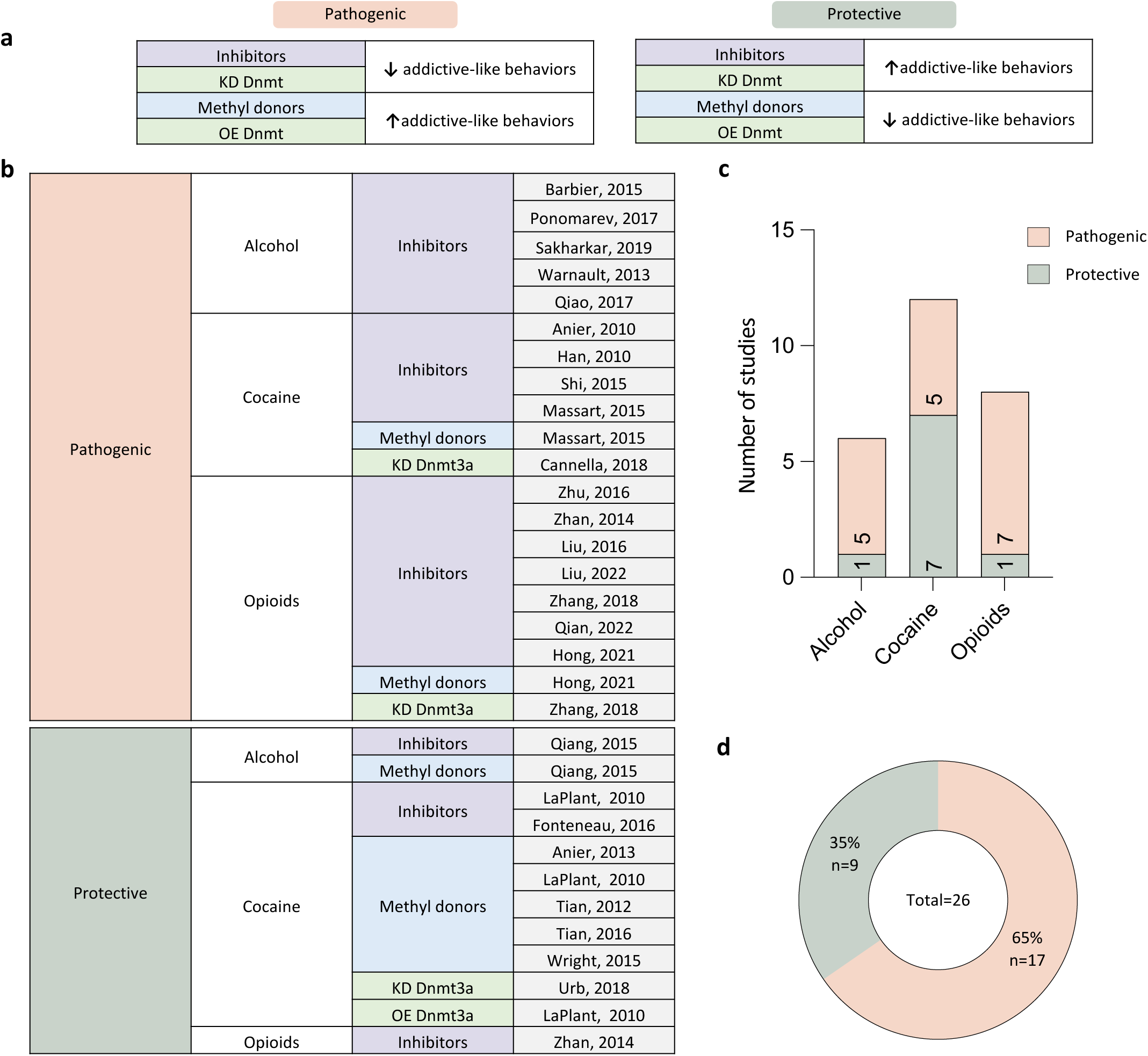
Studies in favor of a pathogenic or protective role of Dnmt enzymes. **a.** To seek for a general interpretation of results detailed in Fig.3, the corresponding studies were classified in 2 groups. The first corresponded to studies that argue for a pathogenic role of Dnmt enzymes, whereby behavioral responses to drugs of abuse were either: (i) blunted upon pharmacological inhibition or genetic knockdown of the enzymes; (ii) or, alternatively, potentiated by dietary supplementation with methyl donors or viral overexpression (OE) of the enzymes. The second group corresponded to studies that argue for a protective role, whereby behavioral effects in opposite directions were observed upon similar experimental manipulations. **b.** Distribution of articles according to this classification. **c-d.** Distribution of studies arguing for a pathogenic or protective role for each drug of abuse (c), or all drugs combined (d).

### 5.1. Alcohol

Six studies evaluated the impact of Dnmt inhibitors (Fig.3b). Three used systemic 5-aza administration: the first found that it blocked alcohol-induced gene expression changes in the VTA, leading to reduced dopaminergic hyperactivity and drug consumption (Ponomarev et al., 2017). The 2 others showed that the expression of Dnmt enzymes was increased in the amygdala (Sakharkar et al., 2019) and NAc (Warnault et al., 2013) during excessive alcohol intake, and that these effects were reversed by 5-aza. In contrast, a single article reported that intracerebroventricular (icv) administration of 5-aza increased chronic intermittent alcohol consumption (Qiang et al., 2014). To go beyond results obtained upon systemic administration, 2 studies performed RG108 or 5-aza local infusion in the medial prefrontal cortex (mPFC). In alcohol-dependent rats, increased DNAm and decreased expression of synaptic plasticity-related genes (such as *Syt2* and *Cacna1a*) were associated with escalated drinking. Interestingly, these effects were reversed by RG108 (Barbier et al., 2015). Similarly, 5-aza decreased alcohol consumption and rescued the downregulation of neurotrophin-3 (Ntf3) induced by alcohol (Qiao et al., 2017). Overall, these results suggest that Dnmt enzymes primarily play a pathogenic role in alcohol molecular and behavioral effects (Fig.4).

### 5.2. Cocaine

Three studies investigated cocaine-induced locomotor sensitization (Fig.3c). Two found that this behavioral response was potentiated by systemic administration of SAM (Anier et al., 2013) and, consistently, reduced by icv infusion of zebularine (Anier et al., 2010). In contrast, the third article showed a reduced response following systemic injection of methionine (Wright et al., 2015). Focusing on another behavior, 2 articles used conditioned place preference (CPP) to explore cocaine reward, and found that it was decreased following methionine injections (Tian et al., 2016, 2012). These contradictory results suggest that manipulating Dnmt enzymes may have bidirectional effects (Fig.4), calling for more precise targeting of specific brain regions, time points and enzymes. Accordingly, local 5-aza infusion in the hippocampus blocked the acquisition but not the retrieval of cocaine CPP, while infusion in the prelimbic cortex was ineffective during acquisition but prevented retrieval (Han et al., 2010). Three additional studies focused on the NAc: a viral shRNA-mediated knockdown of Dnmt3a led to enhanced sensitization (Urb et al., 2020), while RG108 infusion or Dnmt3a overexpression increased or decreased cocaine CPP, respectively (LaPlant et al., 2010; Urb et al., 2020), indicating that Dnmt3a activity may protect against these responses. Interestingly, Tet1 KD or overexpression in the NAc increased or decreased cocaine CPP, respectively, similar to the effects observed for Dnmt3a manipulations (Feng et al., 2015). It may appear counterintuitive that the inhibition of either the *methylation* or *demethylation* pathways may have similar behavioral impact. We note, however, that these enzymes are sensitive to gene dosage (Christian et al., 2020), and that specific effects of each manipulation on levels of 5mC and 5hmC, which have not been characterized at genome-wide scale, may occur at distinct loci and with different kinetics. Along this line, the aforementioned Tet1 KD was sufficient to mimic, in naive mice, the increase in 5hmC levels that the authors had previously identified at 5 candidate genes following cocaine injections. This suggests that, at these loci, the oxidation of 5mC into 5hmC may be less dependent on Tet1 than further oxidations into 5fC and 5caC. More work will be necessary to understand how each Dnmt and Tet enzyme regulates the relative abundance of successive cytosine modifications.

Finally, 5 articles examined voluntary self-administration (SA) of cocaine in rats. One showed that icv infusion of 5-aza or zebularine led to decreased acquisition of operant responding (Feng et al., 2015; Fonteneau et al., 2017). The 4 others focused on relapse, using cue- or drug-induced reinstatement of cocaine-seeking. The first found that systemic methionine reduced cue-induced reinstatement (Wright et al., 2015), in contrast with the 3 other articles that specifically targeted the basolateral amygdala (BLA) or NAc. In the BLA, 5-aza infusion decreased nosepokes during cue- and drug-induced reinstatement (Shi et al., 2015; Wright et al., 2015). In the NAc, shRNA-mediated knockdown of Dnmt3a2 reduced cue-induced reinstatement after 1 and 45 days of withdrawal (Cannella et al., 2018), while RG108 infusion produced a similar decrease after 30 and 60 days of withdrawal, and SAM infusion increased responding (Massart et al., 2015). Overall, results related to protective or pathogenic roles of DNAm enzymes were comparatively more heterogeneous for cocaine than for alcohol (Fig.4).

### 5.3. Opioids

Eight articles investigated the impact of 5-aza (Fig.3d). The first showed that intrahippocampal infusion prevented morphine-induced DNA hypermethylation at the glucocorticoid receptor locus (GR, a key component of the stress axis dysregulated in SUD), an effect associated with reduced physical signs of (Zhu et al., 2017). Three additional studies from the same group used CPP and conditioned place aversion (CPA). For CPP, dissociations were observed depending on the timing and brain region targeted for 5-aza infusion: in the hippocampus (CA1), a reduction in the acquisition and consolidation of morphine CPP was observed; in the prelimbic cortex (PL), retrieval was potentiated (Zhang et al., 2014); in the BLA, reconsolidation was impaired (Liu et al., 2022). For naloxone-induced CPA, 5-aza infusions in the agranular insula (AI) or BLA disrupted reconsolidation (Liu et al., 2016).

The remaining studies focused on opioid SA in rats. Three used 5-aza infusions in the BLA, CA1, and NAc, and observed reduced drug-seeking (Hong et al., 2020; Qian et al., 2022; Zhang et al., 2020). In the CA1 region, a knockdown of Dnmt3a produced a similar effect (Zhang et al., 2020). Consistently, systemic administration of L-methionine, a methyl donor, had the opposite effect and increased drug-seeking (Hong et al., 2020). A single article explored the function of Tet enzymes in opioid SA. In the CA1, Tet1 and Tet3 were dysregulated with distinct kinetics: after 1 day of morphine SA, Tet3 expression was increased; after 7 days, Tet3 returned to baseline while Tet1 was downregulated. Interestingly, Tet3 knockdown accelerated morphine SA, while overexpression of Tet1 attenuated morphine SA, suggesting that the activity of the 2 enzymes was protective against the acquisition of this behavior (Jiang et al., 2021).

Altogether, a majority of the studies described above indicate that alcohol, cocaine or opioids recruit Dnmt or Tet enzymes, which in turn contribute to pathological SUD-related behaviors. A minority of reports also suggests that in certain experimental contexts, these enzymes may act to counteract such behaviors and rather represent a protective mechanism. Importantly, the specific enzymes that may be recruited by each drug of abuse across different brain regions or cell types, and the loci at which they act (beyond candidate genes), remain to be more systematically assessed.

## 6. Drug-induced DNA methylation changes

To investigate methylomic effects of drugs of abuse, 3 approaches have been used. The first corresponds to the interrogation of global DNAm levels to generate a single measure for each biological sample, using methods such as ELISA. A more precise approach consists in assessing candidate genes, most frequently their promoters. Finally, recent progress now allows mapping drug-induced changes in an unbiased manner, focusing on DNA regions rich in CG sites, or the full methylome.

### 6.1 Global measures

Following alcohol exposure (Supplementary Figure1), DNAm levels were increased in the PFC, but not affected in the mouse amygdala or hippocampus (Yang et al., 2021). In the rat, an increase was reported in the NAc (Niinep et al., 2021), while another study also found increases in the PFC and NAc, with no changes in the amygdala or hippocampus (Barbier et al.,2015). For other drugs of abuse, results were more heterogeneous. For cocaine, no changes were detected in the whole mouse brain (Chao et al., 2014; Fragou et al.,2013) or in the corpus callosum (Nielsen et al., 2012). In the NAc, no changes were observed in mice (Feng et al., 2015; Tian et al., 2012), while either a decrease (Wright et al., 2015) or an increase (Anier et al., 2018) were found in rats. A single study in the hippocampus found an increase in 5hmC following abstinence (Sadakierska-Chudy et al.,2017). In the PFC, a decrease in mice (Tian et al., 2012) but an increase in rats (Saad et al., 2021) were reported, with the later study also describing an increase in the caudate putamen (Saad et al., 2021). For opioids, no significant effects were detected in the mouse NAc or PFC after morphine (Tian et al., 2012). Other studies found significant decreases: in the striatum, after morphine injections (Joanna et al., 2017); in the hippocampus, following heroin SA (Chen et al., 2019) or oxycodone injections (Fan et al., 2019); in the VTA, after oxycodone injections (Fan et al., 2019). In contrast, a selective 5hmC increase following oxycodone injections was described in the hippocampus (Fan et al., 2021). For nicotine, increases were reported by 2 rat studies in 3 regions (NAc, orbitofrontal and prefrontal cortices (Mychasiuk et al., 2013; Nguyen et al., 2019), while a mouse study reported a decrease in 2 regions (cortex and striatum), and no effects in the hippocampus (Buck et al., 2019). Finally, regarding amphetamine, 2 studies found increases in the rat NAc and PFC (Mychasiuk et al., 2013; González et al., 2018). While it is possible that part of these discrepancies may reflect methodological (time-point, administration route) or biological (region, drug doses, species) differences, such measures of global DNAm levels should be considered with caution. As mentioned above, they reflect highly variable patterns of DNAm among millions of CG and non-CG sites. Rather than their average, it is their discrete cell type- and locus-specific distribution that mediates changes in gene expression. This is illustrated by results from Feng et al (Feng et al., 2015): although cocaine did not induce any global differences in genome-wide levels of 5mC or 5hmC, local changes in 5hmC (both up or down) were nevertheless detectable at 11,500 loci. Therefore, investigating DNAm at higher genomic and cellular resolution will be necessary to advance SUD research.

### 6.2. Candidate gene studies

#### 6.2.1. Alcohol

Twelve articles investigated 15 genes across 12 regions. Most articles focused on a single region, and only 4 considered more than one. Since no individual gene was investigated by more than 1 article in any given region, no findings were replicated (Fig.5). Six articles investigated DNAm only (i.e. with no consideration of gene expression), among which 4 showed negative results, with no evidence for alcohol-induced DNAm changes in the promoters of *Pdyn* and *Pnoc* in the amygdala (D’Addario et al., 2012), of *Crhr1* and *Fkbp5* in the hypothalamus (Todkar et al., 2015), of *Drd2* in the NAc (Feltmann et al., 2018), as well as in the promoter and first intron of *Maoa* in the NAc and dorsal striatum (DS; Bendre et al., 2019). In contrast, alcohol led to reductions in DNAm in the promoter or at the transcription start site (TSS) of *Hif3a* and *Slc10a6* in the amygdala (Krishnan et al., 2022), and along 7 CGI of *Bdnf* in the hippocampus (Stragier et al., 2015).

**Figure 5.**
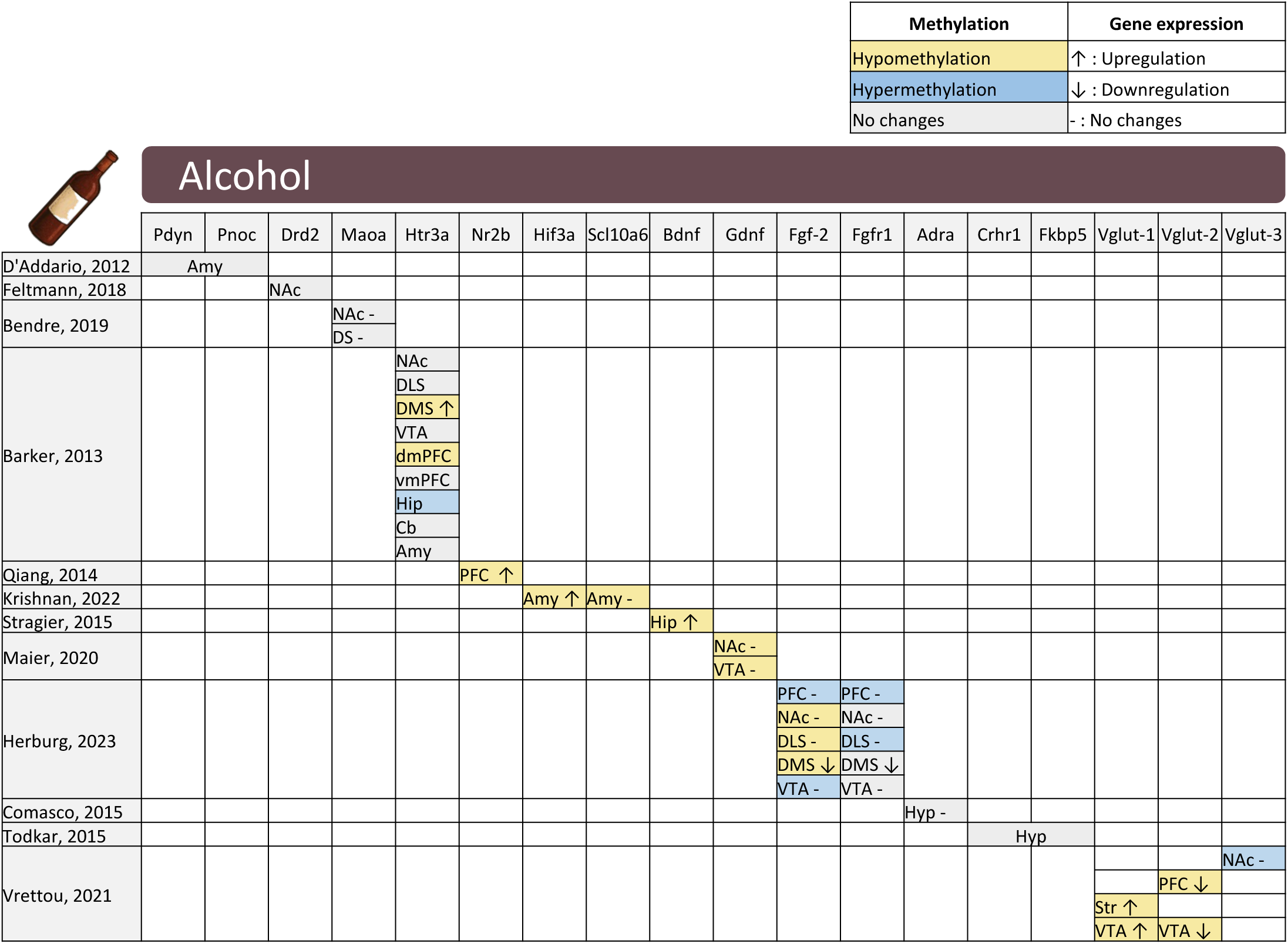
Candidate gene studies on alcohol. The table summarizes the brain region and genes that were investigated. The over-representation of empty cells illustrates the fact that no specific gene was investigated by more than a single study in any brain region. Yellow and blue indicate a decrease or an increase in DNAm following alcohol exposure or consumption, respectively, while grey indicates no significant changes. Upward or downward arrows indicate an increase or a decrease in gene expression, respectively, while “-” indicates no significant changes. Similar tables for cocaine, opioids, and methamphetamine are available as Supplementary Figures 2-4. Abbreviations: Amy, amygdala; Cb, cerebellum; DLS, dorsolateral striatum; DMS, dorsomedial striatum; DS, dorsal striatum; Hip, hippocampus; Hyp, hypothalamus; NAc, nucleus accumbens; PFC, prefrontal cortex; dmPFC, dorsomedial prefrontal cortex; vmPFC, ventromedial prefrontal cortex; Str, striatum; VTA, ventral tegmental area.

The 6 remaining articles analyzed both DNAm and gene expression. One found no changes in promoter DNAm or in the expression of *Adra2a* in the hypothalamus (Comasco et al., 2015). In contrast, chronic intermittent exposure decreased DNAm in the *Nr2b* promoter in the mouse PFC, which correlated with increased expression of this glutamatergic receptor subunit (Qiang et al., 2014). The last 3 studies investigated multiple brain regions. DNAm was decreased in the promoter of *Gdnf* in the NAc and VTA of alcohol-treated rats, although its expression was unaffected (Maier et al., 2020). Bidirectional changes in DNAm were found in the *Fgf-2* promoter across 6 regions, but gene expression was affected in the dorsomedial striatum only (DMS, (Herburg et al., 2023)). The same study also found minor effects for another gene, *Fgf1*, with slight DNAm differences at individual CG sites and decreased expression, again only in the DMS. In another rat study, *Vglut1, 2* and *3* were investigated in 4 regions (NAc, PFC, striatum and VTA). Results showed subtle gene and region-specific variations, with low correlation between DNAm and mRNA (Vrettou et al., 2021). Finally, the *Htr3a* gene promoter was investigated in 9 regions (Barker et al., 2013). Alcohol led to increased DNAm in the hippocampus, and a decrease in the dorsomedial PFC and DMS, but not the other 6 regions. This was associated with increased expression in the DMS, although other regions were not investigated.

#### 6.2.2. Cocaine

Twelve articles investigated 16 genes across 8 brain regions (Supplementary Figure2). As with alcohol, most of them (n=9) investigated a single region. Two examined DNAm only, using SA models. The first reported that DNAm at the *FosB* and *Crem* promoters increased in the hippocampus but decreased in the PFC following extended cocaine access (Ajonijebu et al., 2017). The other focused on the corpus callosum and genes related to oligodendrocytes, and showed that, after 30 days of withdrawal, the *Sox10* promoter was hypomethylated, while no changes were observed for *Plp1* and *Mbd* (Nielsen et al., 2012).

The remaining 10 articles examined both DNAm and gene expression. One study analyzed whole-brain microglia and found increased methylation of the miR-124 promoter, accompanied by decreased expression (Guo et al., 2016). Two investigated the PFC and reported decreased promoter methylation of *Homer2* (Ploense et al., 2018) and *Hcrtr1* (also known as *Ox1r*) (Saad et al., 2019), both associated with increased gene expression. The remaining 7 articles investigated the striatum. One reported that 45 days of cocaine withdrawal led to *Glt-1* promoter hypomethylation and increased *Glt-1* expression in the NAc (Kim et al., 2018). Another examined 3 genes and found promoter hypermethylation and downregulation of *Pp1c* and *A2aR*, along with promoter hypomethylation and upregulation of *Taarb7*, in both the NAc and the LHb (Vaher et al., 2020). Two studies further focused on *Pp1c* to examine how MeCP2 may contribute to these correlations. They reported consistent findings, as cocaine increased the binding of MeCP2 to the *Pp1c* promoter in both the CPu and NAc (along with reduced gene expression), in agreement with its repressive function (Pol Bodetto et al., 2013; Anier et al. 2010). Using a similar approach, another study found increased MeCP2 binding at the *Cdkl5* promoter and decreased expression in the striatum (Carouge et al., 2010). Beyond MBD proteins, DNAm also regulates the binding of transcription factors (TF). In the NAc, cocaine CPP decreased DNAm at the promoter of the *Bdnf IV* splice variant, which was associated with increased binding of the *Myb* TF, and increased transcript expression (Tian et al., 2016). This mechanism at the *Bdnf* promoter was documented by an *in vitro* Luciferase assay. In parallel, behavioral and molecular effects of cocaine were prevented by the methyl donor methionine. Finally, in addition to the modulation of MBD and TF binding in *cis*, drug-induced changes in DNAm may also act by modifying the 3D organization of the genome. This was explored in an interesting study that focused on a 1.5 Mb loop formed between the *Auts2* and *Caln1* genes. Upon cocaine injections, increased DNAm at the genomic sites where the 2 loci interact led to an opening of the loop, which induced an accumulation of the activating H3K4me3 histone modification and increased expression of both genes (Engmann et al., 2017), specifically in medium spiny neurons expressing the *Drd2* receptor.

#### 6.2.3. Opioids

Six articles investigated 10 genes across 12 regions, with only one study that considered multiple regions (Supplementary Figure3). Half of these studies investigated DNAm only. The first one found that rats exposed to morphine did not show any DNAm differences in the promoter of *Drd2* (Rodrigues et al., 2012). In a second study, a single CG was differentially methylated in the *Crhbp* promoter following heroin exposure (McFalls et al., 2016). The third one assessed 6 genes (*Bdnf*, *Comt*, *Il1b*, *Il6*, *Nr3c1*, and *Tnf)* in 10 regions of the rat brain, after acute or chronic morphine. Results showed that DNAm varied at a few promoters only, and that these changes were variable across genes, duration of exposure and regions (Barrow et al., 2017).

In the articles that examined both gene expression and DNAm, hypomethylation in the *Egr2* promoter, along with decreased gene expression, were observed in the mPFC following heroin SA (Imperio et al,. 2018). Another study reported in the hippocampus an hypermethylation at the promoter of the *Nr3c1* 17 variant (which encodes GR), and increased gene expression, following chronic morphine (Zhu et al., 2017). Interestingly, this article also showed that morphine led to increased expression and binding at the *Nr3c1* promoter of the Dnmt1 and Mbd2 proteins, likely accounting for the observed changes. The last study showed that *Dnmt3a* expression was decreased in the hippocampus following heroin SA, an effect driven by an increase in Ube2b-mediated ubiquitination and degradation of the enzyme (Chen et al., 2019). Through elegant in vivo experiments, this decrease in Dnmt3a activity was shown to lead to reduced promoter DNAm and increased expression of *Camkk1,* which in turn drove actin polymerization and contributed to drug-seeking.

#### 6.2.4. Methamphetamine

Nine articles assessed 15 genes across 5 regions, among which only 4 examined more than one region (Supplementary Figure4). All explored DNAm in combination with gene expression. In the striatum, decreased DNAm was reported at the promoters of the *Scna* (Biagioni et al,. 2019) and *Glua2* (Jayanthi et al.,2014) genes, while their expression was increased and decreased, respectively. In the NAc, the same group also found increased expression of 3 potassium channels (*Kcna1*, *Kcna3* and *Kcnn1*) following 30 days of abstinence, along with decreased promoter DNAm (Jayanthi et al.,2019). Another group examined across 2 brain regions the impact of methamphetamine on 2 types of repetitive elements, Long Interspersed Elements (LINE-1) (Moszczynska et al., 2015) and Short Interspersed Elements (SINEs) (Moszczynska et al., 2017). In the dentate gyrus, a decrease in DNAm at a single CG was observed for *Pabp1*, a SINEs regulatory protein, 1 hour after binge methamphetamine, but not at later time points (Moszczynska et al. 2017). In the striatum, DNAm of LINE-1 elements was increased 7 days after methamphetamine binge injections, which was associated with decreased expression of ORF-1, the protein they encode (Moszczynska et al.,2015). Two other articles investigated different CGI of *Bdnf IV* in the PFC and hippocampus. In the hippocampus, *Bdnf IV* was hypomethylated and upregulated. In the PFC, either increased DNAm with no expression changes (Iamjan et al.,2021) or decreased DNAm and upregulation were observed (Salehzadeh et al,. 2020), possibly reflecting locus-specific regulation. Finally, 2 studies analyzed genes related to synaptic plasticity in the PFC and hippocampus. In the PFC, methamphetamine reduced the expression of *Arc*, *Egr2*, *Fos*, *Nr4a1* and *Syn*, with divergent methylation patterns: no changes for *Egr2* and *Nr4a1*, hypomethylation for *Arc*, and hypermethylation for *Fos* (Cheng et al,. 2015) and *Syn* (Fan et al.,2020). In contrast, in the hippocampus, the same genes were also downregulated but with reduced methylation at the *Nr4a1* promoter, no changes for *Arc*, *Egr2* and *Fos*, while *Syn* was upregulated and hypomethylated (Cheng et al,. 2015; Fan et al.,2020). These results again highlight a poor correlation between drug-induced changes in gene expression and DNAm, at least when individual genes are considered.

Altogether, these candidate gene studies conducted at bulk tissue level indicate a loose relationship between promoter DNAm and gene expression. In addition, dissociations among studies that investigated multiple regions suggest that the expression of individual genes may be regulated by region-specific mechanisms.

### 6.3. Genome-wide studies

The candidate studies described above collectively indicate that exposure to drugs of abuse modulate DNAm at specific loci. By nature, these approaches face a risk of confirmation bias and cannot identify new genes associated with SUD. Another limitation is that most of them analyzed promoters, where changes in DNAm have been primarily characterized in relation to large changes in gene expression, such as during cellular differentiation. In post-mitotic neurons, the modest expression changes induced by drugs of abuse more likely rely on subtle regulatory mechanisms at tissue- and cell-type specific enhancers, transcription factor binding sites (Kreibich and Krebs 2023), or intron/exon junctions regulating splicing (Lev Maor et al., 2015). As such, hypothesis-free and pan-genomic analyses are necessary to go beyond promoters and candidate genes. These efforts are summarized below, depending on whether a genome-wide analysis of gene expression was followed by investigation of DNAm at a few genes, or vice versa.

#### 6.3.1. Transcriptome first

##### 6.3.1.1. Alcohol

The aforementioned article that investigated global DNAm and Dnmt inhibition following alcohol abstinence also performed a transcriptomic analysis in the mPF, using RNA-sequencing ((Barbier et al., 2015), see sections 5.1 and 6.1). Results indicated that 784 genes were nominally differentially expressed, and enriched in GO terms related to transcription and neurotransmission. The downregulation of 7 genes was confirmed by qPCR and, for 4 of them, shown to be prevented by RG108. In particular, the downregulation of *Syt2* (a calcium sensor mutated in myasthenic syndrome) was associated with hypermethylation of a single CG site in its first exon, an effect blocked by RG108. Finally, a viral shRNA KD of *Syt2* led to higher alcohol consumption when the drug was paired with quinine (adulteration), uncovering a role for that gene in compulsive-like intake.

##### 6.3.1.2. Cocaine

Microarrays were used to analyze the effects of SAM and chronic cocaine, alone or in combination, in the mouse NAc (Anier et al., 2013). Surprisingly, SAM pretreatment blunted roughly half of cocaine-induced adaptations, whether they corresponded to up- or down-regulations. DNAm was then investigated at CGI within promoters of 3 genes with most significant differential expression, using methylated DNA immuno-precipitation followed by qPCR (MeDIP-qPCR). For 2 of these, cocaine-induced changes in DNAm were partially blocked by SAM pretreatment. Based on further in vitro and in vivo work, the authors proposed that such modulatory effects of SAM may be partly mediated by decreased Dnmt3a expression.

##### 6.3.1.3. Opioids

RNA-sequencing was used to analyze the mPFC in a rat model of the progressive devaluation of a natural reward (saccharine) induced by heroin SA (McFalls et al., 2022). Two groups of rats were distinguished based on whether the suppression of saccharine intake (i.e., its devaluation) was large or small, and compared to rats self-administering heroin without any access to saccharin. Sixty genes were differentially expressed across those groups, among which 10 were investigated by qPCR, and 6 validated. Using bisulfite amplicon sequencing, changes in DNAm at 6 individual CG sites were identified at CGI located among 5 of these genes. Of note, LINE elements were also explored, but no DNAm changes observed, contrasting with aforementioned methamphetamine studies (Moszczynska et al., 2015; Moszczynska et al., 2017 ; Cadet et al., 2016).

##### 6.3.1.4 Methamphetamine

Microarrays were used to identify 503 genes that were differentially expressed in the rat NAc following a single methamphetamine injection (Jayanthi et al., 2018). The authors then focused on increased expression of 2 stress-related genes, *Crh* and *Avp*. MeDIP-qPCR showed lower 5mC and higher 5hmC at the *Crh* promoter and *Avp* gene body, respectively, while ChIP-qPCR documented higher binding of the Tet1 and Tet3 enzymes at these loci.

#### 6.3.2. DNA methylation first

Genome-wide DNAm analyses have been performed in 10 studies (Fig.6) following exposure to cocaine (n=6), opioids (n=2), alcohol (n=1), or methamphetamine (n=1). Each study investigated a single brain region. Eight studies used immunoprecipitation of methylated (n=2; Fig.6a) or hydroxymethylated (n=2) DNA, or immunoprecipitation of MBD proteins (n=3), followed by sequencing (MeDIP-seq, hMeDIP-seq and MBD-seq, respectively), thereby generating peak-like, low resolution measures of DNAm. More recent work used bisulfite conversion and sequenced either the Whole Genome (WGBS) or a Reduced Representation (RRBS), providing methylomic measures at single-base resolution (n=3). Of note, only 2 studies focused specifically on 5hmC (Feng et al., 2015; Cadet et al., 2016), while others used approaches that did not differentiate between 5mC and 5hmC, indicating that the respective roles of these modifications remain poorly defined.

**Figure 6.**
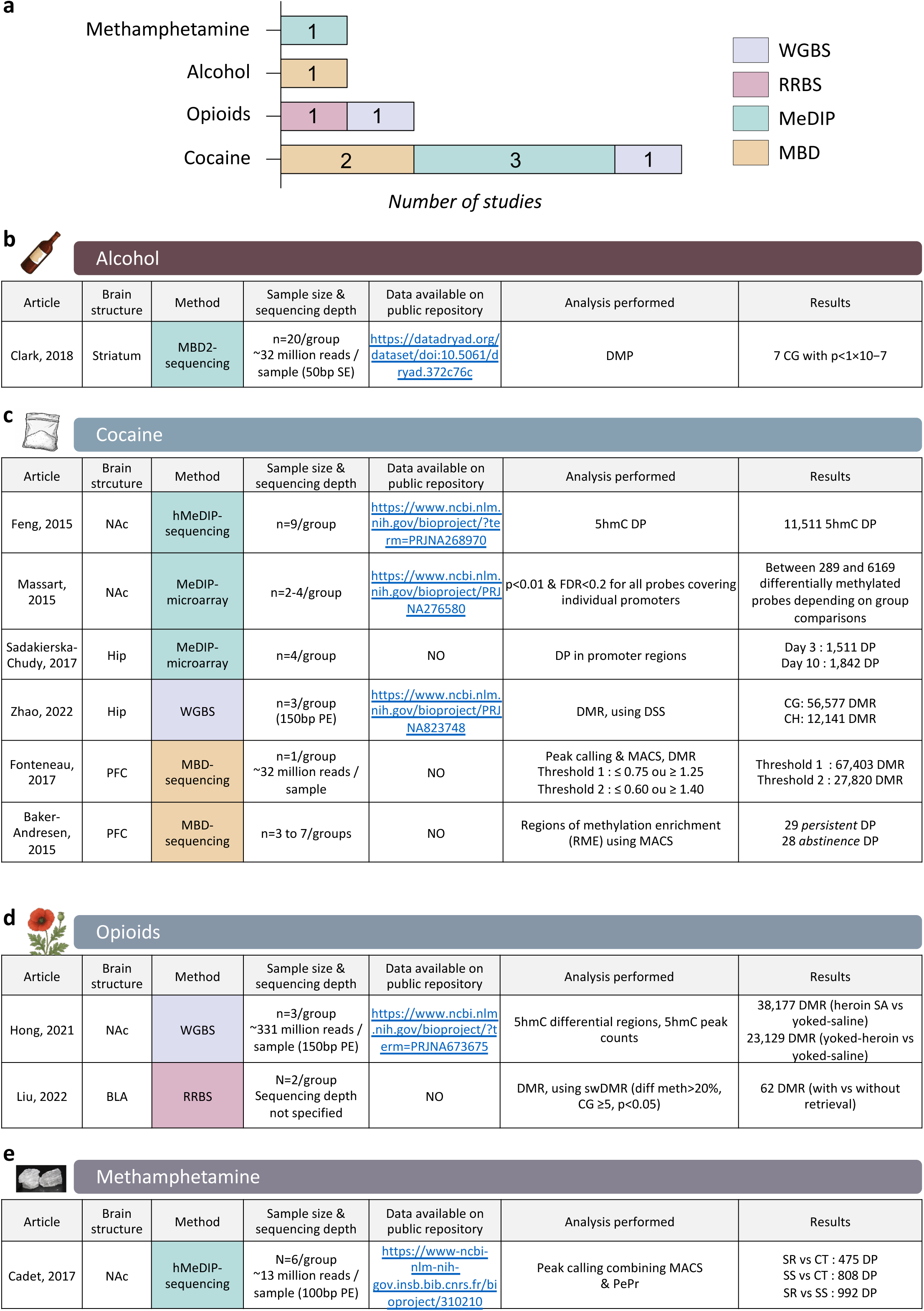
Genome-wide studies of DNA methylation in rodents exposed to drugs of abuse in adulthood. **a.** Distribution of types of studies across drugs. **b-e.** Tables summarizing studies conducted on alcohol (b), cocaine (b), opioids (d) or methamphetamine (e). Abbreviations: BLA, basolateral amygdala; bp, base pairs; CT, control; DMP, differentially methylated positions; DMR, differentially methylated regions; DP, differential peaks; FDR, false discovery rate; Hip, hippocampus; MBD2-sequencing, immunoprecipitation of the MBD2 protein coupled with sequencing; MeDIP- and hMeDIP-microarray, immunoprecipitation of methylated or hydroxymethylated DNA coupled with microarrays; NAc, nucleus accumbens; PE, paired-end reads; PFC, prefrontal cortex; RRBS, reduced representation bisulfite sequencing; SR, shock-resistant; SS, shock-sensitive; WGBS, whole-genome bisulfite sequencing.

##### 6.3.2.1 Alcohol

Acute changes in DNAm were characterized in mice following a single systemic injection of alcohol (2g/kg), using MBD2-sequencing and a large sample size (n=20/group (Clark et al., 2018)). This enabled the analysis of 22 million CG in both the brain (striatum) and blood, with the aim of identifying peripheral biomarkers. Individual CG showed alcohol-induced changes in each tissue (Fig.6b), and a significant overlap was detected among top results from both tissues (at a low, 20-kb resolution).

##### 6.3.2.2 Cocaine

The effects of cocaine on DNAm were investigated by 6 studies (Fig.6c). In the NAc, the first study used MeDIP and microarrays to analyze promoters in a rat SA model (Massart et al.,2015). Four experimental groups were investigated following either 1 or 30 days of cocaine withdrawal, with or without exposure, at each time-point, to a session of cue-induced reinstatement. Interestingly, contrasting with the classical view of DNAm as a stable epigenetic mark, each 1-hour session of reinstatement was sufficient to trigger hundreds of DNAm changes. In addition, a higher number of differences were observed after 30 than after 1 day of withdrawal, consistent with the behavioral incubation of drug-seeking. Gene expression changes were also characterized, and showed modest but statistically significant overlap with DNAm changes, again highlighting a complex relationship. The second NAc study (already mentioned in section 6.1) used a method based on chemical labelling and affinity enrichment of 5hmC (Feng et al., 2015). The in-depth analyses conducted by the authors showed that bidirectional changes in 5hmC were detectable at 11,500 loci. These changes affected genes involved in neuronal plasticity and chromatin organization, occurred notably at exon boundaries and enhancers (defined by the presence of H3K27a and H3K4me1, but absence of H3K4me3), and significantly overlapped with genes showing drug-induced changes in expression or alternative splicing.

In the hippocampus, the first study used MeDIP and a promoter array, as well as gene expression microarrays, to investigate a rat model of SA (Sadakierska-Chudy et al.,2017). At the transcriptomic level, 26 genes were found upregulated after both 3 or 10 days of daily extinction training during drug withdrawal. For DNAm, 1,511 and 1,842 differential peaks (DP) were identified at each time-point, although no global comparisons of results from the 2 omic layers or GO analysis were conducted. The second study focused on the role of cocaine as a worsening factor in a mouse model of cognitive deficits induced by HIV infection (Zhao et al., 2022). DNAm was analyzed using WGBS, enabling the identification of successive CG sites showing changes in a similar direction across groups, defined as differentially methylated regions (DMR). Results indicated that both HIV infection (modelled using a transgenic line overexpressing the viral protein Tat) and chronic cocaine, as well as their interaction, associated with large numbers of DMR (>30,000 each) broadly distributed across all chromosomes, and significantly associated with differentially expressed genes. GO analyses of DMR pointed to alterations of neuronal function and the extra-cellular matrix. At histological level, these effects were associated with higher spine density, suggesting that cocaine-induced changes led to synaptic dysfunction.

In the PFC, the first study used MBD-seq to compare 3 groups of rat self-administering cocaine (with or without 5-aza infusion) or a saline solution, as control (Fonteneau et al., 2017). Because of a low sample size (n=1), DP were defined using thresholds on methylation ratios. Counterintuitively, 3 times more DP corresponding to increased rather than decreased DNAm were observed in the Cocaine+5-aza sample compared to the Cocaine one. Around half of these DP were located in gene bodies, 9% in promoters, and the remaining in intergenic regions. Ingenuity Pathways Analysis suggested that 5-aza treatment increased cocaine intake by modulating DNAm at genes related to axonal growth and synaptogenesis. The second study also used MBD-seq and a SA model, but in mice (Baker-Andresen et al., 2015). DNAm was analyzed after 1 or 21 days of drug withdrawal following a cue-induced reinstatement session, as well as in 2 control groups of yoked or drug-naive mice. Interestingly, this study specifically analyzed the methylome of PFC neurons (using flow cytometry to isolate nuclei expressing the NeuN neuronal maker, an approach that has since been frequently used). Few significant results were identified, with 29 *persistent* DP (detected at both 1 and 21 days of withdrawal) and 28 *abstinence-associated* DP (emerging specifically after 21 days). Some of these were associated with changes in the expression of nearby genes and, in some cases (*Cpeb4, Mctp1*), in the expression of specific isoforms, reflecting a potential role of DNAm in regulating drug-induced alternative splicing.

##### 6.3.2.3 Opioids

Two opioid studies were conducted in rats. The first used WGBS to analyze heroin SA in the NAc (Hong et al., 2020): 38,177 DMRs were identified when comparing the heroin SA and yoked-saline groups, and 23,129 DMRs between the yoked-heroin and yoked-saline groups. Surprisingly, the genomic distribution or GO enrichment of these numerous findings was not investigated. Rather, the authors focused on 2 genes for validation of DMR and their transcriptional impact (*Gabrd* and *Gabrp*), and then conducted global manipulation of DNAm using 5-aza and methionine (see section 5.3). The second study also performed Dnmt inhibition (section 5.3) to demonstrate a role for the BLA during retrieval of morphine CPP (Liu et al., 2022). RRBS was used to identify 62 DMR associated with retrieval. Among the 13 DMR located in promoters, 3 affecting neural plasticity-related genes were prioritized for validation, resulting in a positive finding for *Gnas*, which also exhibited decreased expression. Again, the GO enrichment of DMR, or the biological relevance of those located outside promoters, were not addressed.

##### 6.3.2.4 Methamphetamine

One study investigated rats that self-administered methamphetamine, which were separated in 2 groups depending on whether operant responding decreased when it was paired with electric footshocks (Cadet et al., 2016). 5hmC was investigated using hMeDIP-seq and compared across shock-resistant (SR) and shock-sensitive (SS) rats. A large number of differentially hydroxymethylated peaks were detected, mostly in repeated elements and intergenic regions. Using Ingenuity Pathway Analysis, these peaks were found enriched in genes related to cognition, synaptic plasticity and potassium transport. Among the latter group, 5hmC differences were associated with modest changes in expression.

Overall, these genome-wide studies showed major differences in sample size, analytical method and significance threshold (Fig.6), resulting in vastly different numbers of findings. Raw data or code were frequently not accessible, limiting the potential for reuse and meta-analyses. Nevertheless, 2 main conclusions emerge: first, the epigenetic reprogramming recruited by drugs of abuse is widespread throughout the genome, and largely exceeds adaptations previously uncovered at candidate genes; second, most changes occur outside promoters (see below and Fig.7).

**Figure 7.**
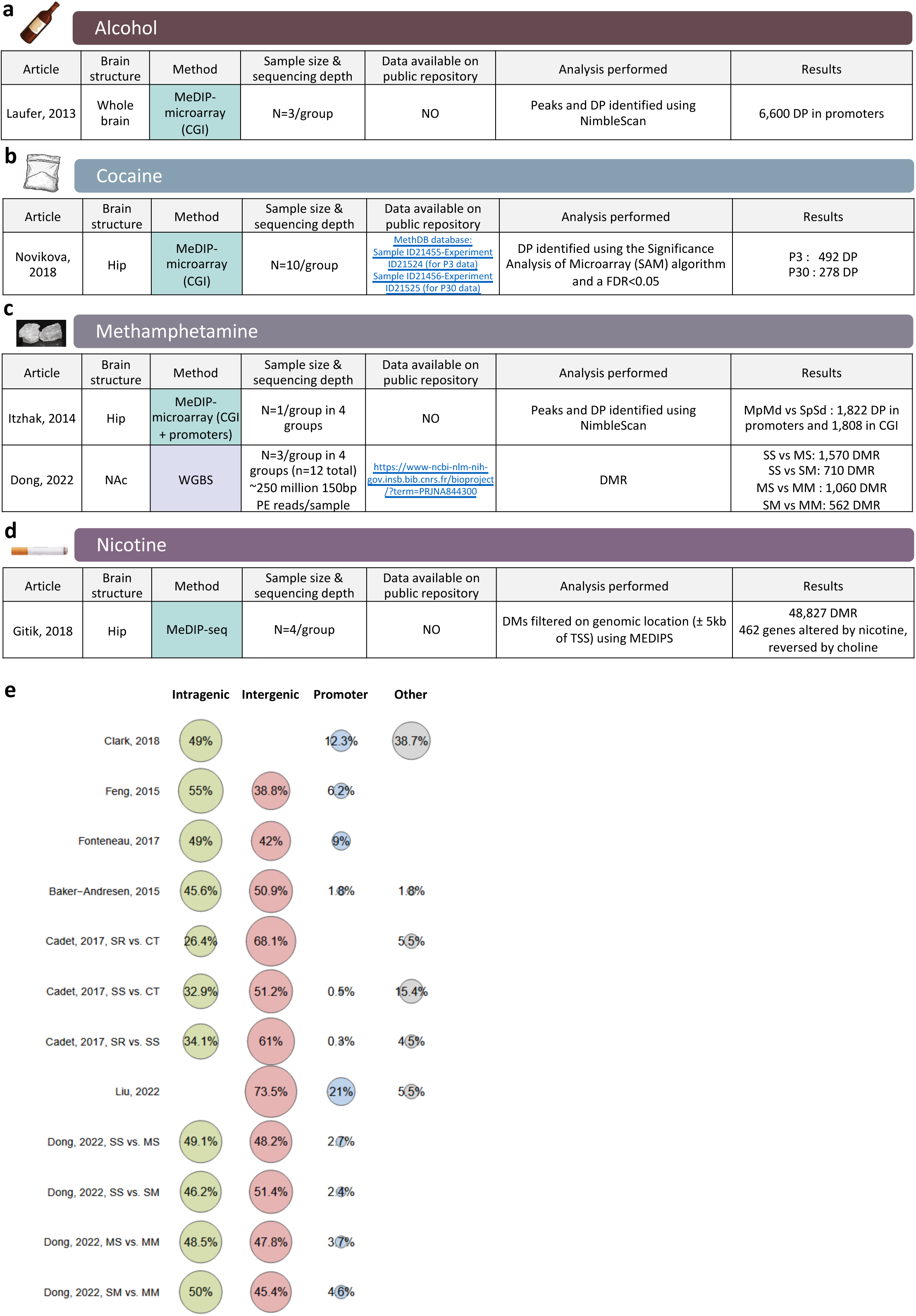
Genome-wide studies in rodents exposed to drugs of abuse during gestation or adolescence. **a-d.** Tables summarizing studies conducted on alcohol (a), cocaine (b), methamphetamine (c), or nicotine (d). **e.** Genomic distribution of differences observed as a function of exposure to drugs of abuse in adulthood (detailed in Fig.6) or during gestation and adolescence (panels a-d from this figure). “Other” indicates the proportion of findings for which genomic localization was not detailed. Itzhak et al compared 4 groups, including: SpSd, saline-exposed pups reared by biological saline-treated dams; MpMd, methamphetamine-exposed pups reared by biological methamphetamine-treated dams (Itzhak et al., 2015). Dong et al compared 4 groups of mice exposed to: saline injections during gestation and adulthood (SS); saline during gestation and methamphetamine in adulthood (SM); methamphetamine during gestation and saline in adulthood (MS); methamphetamine during gestation and in adulthood (MM; (Dong et al., 2022)). Main results are summarized here (see original publications for additional group comparisons). Abbreviations: CGI, CG islands; DP, differential peaks; DMR, differentially methylated regions; Hip, hippocampus; MeDIP-microarray, immunoprecipitation of methylated DNA coupled with microarrays; NA, not available; NAc, nucleus accumbens; P3 and P30, post-natal days 3 and 30.

## 7. Developmental effects of drugs of abuse on DNA methylation

While aforementioned studies investigated adult animals, early-life exposure to drugs of abuse during gestation or adolescence may have long-lasting epigenetic impact, which may contribute to the risk of SUD in adulthood.

### 7.1. Drug exposure during adolescence

Three studies investigated in the mouse the long-term methylomic effects of drug exposure during adolescence: 2 on alcohol, 1 on nicotine. For alcohol, both articles focused on candidate genes. The first showed that intermittent exposure from post-natal day 25 to 55 (P25-55) led to increased DNAm at the promoter of the choline acetyltransferase *Chat* gene in the basal forebrain of adult mice (Vetreno et al., 2020). Using the same model, the second study reported increased global activity of Dnmt enzymes in the amygdala, as well as local increases in 5mC (but not 5-hmC) at the *Bdnf* and *Npy* promoters (Sakharkar et al., 2019). Interestingly, these effects were reversed by 5-aza treatment. For nicotine, the effects of adolescent exposure on adult hippocampal DNAm were investigated using MeDIP-sequencing (Gitik et al., 2018). While genome-wide data were generated, the authors focused on promoters and prioritized genes for which nicotine-induced DNAm changes were reversed by supplementation with choline, a nutrient with numerous functions (acetylcholine and phospholipid precursor, methyl donor). The resulting 462 genes were enriched in chromatin remodeling factors, suggesting an epigenetic contribution to long-term learning deficits induced by nicotine adolescent exposure.

### 7.2. Drug exposure during gestation

Ten articles assessed how maternal drug exposure during gestation affected DNAm in offspring: 5 on alcohol, 2 on methamphetamine, 2 on nicotine, 1 on cocaine. For alcohol, the first study reported that the adult offspring of dams exposed to alcohol drinking during pregnancy showed a decrease in global DNAm, an effect prevented by choline supplementation (Bottom et al., 2020). The same group also investigated candidate genes, and reported increased promoter DNAm and decreased expression of the *Drd2* dopamine receptor in the pituitary gland (Gangisetty et al., 2015), as well as similar changes affecting *Pomc* and *Sry* in the hypothalamus (Gangisetty et al., 2021). A third study focused on *Pdyn*, and found higher expression and decreased DNAm at a single CG in the promoter (Wille-Bille et al., 2020). Finally, one study used a genome-wide approach (MeDIP coupled with an array targeting CGI) to investigate whole brain tissue (Laufer et al., 2013). More than 6,600 promoters were identified as differentially methylated, with an enrichment for *Cdk5* and *Pten* signalling pathways, as well as for binding sites for CTCF, a protein implicated in chromatin organization, which may therefore contribute to long-term consequences of fetal alcohol exposure.

For cocaine, a single study conducted in the hippocampus used MeDIP coupled with a CGI array to explore how maternal exposure impacted DNAm at P3 and P30 in the offspring (Novikova et al., 2008). At P3, cocaine-induced DNAm changes were observed at 492 CGI. Interestingly, a significant proportion of these sites (67%, n=186) were still affected at P30, although frequently in an opposite direction (43%, n=80), suggesting complex kinetics in drug-induced epigenetic dysregulation.

For methamphetamine, 2 studies used genome-wide approaches. The first investigated the hippocampus using MeDIP coupled with an array covering CGI and promoters (Itzhak et al., 2015). In addition to examining the long-term effects of *in utero* drug exposure, the authors explored the influence of maternal care from dams exposed or not to methamphetamine, using a cross-fostering design. Interestingly, comparable numbers of promoters were differentially methylated depending on whether drug exposure occurred in the offspring in utero or in the dam before and during pregnancy. These findings suggest that early-life behavioral experiences, such as maternal care, can substantially contribute to the intergenerational transmission of drug-related phenotypes. The second methamphetamine study focused on the NAc using WGBS (Dong et al., 2022). The authors compared 4 groups of mice corresponding to the offspring of dams exposed to the drug before and during pregnancy and/or in adulthood. Results indicated that differential methylation at individual CG sites was more strongly associated with maternal than with adult exposure, consistent with the notion that brain development represents a period of heightened vulnerability.

Finally, 2 studies focused on nicotine and explored global DNAm. One examined the striatum (Ilott et al., 2014), the other the striatum and frontal cortex (Buck et al., 2019). Results were contradictory in the striatum, with either no changes (Ilott et al., 2014) or a global hypomethylation (Buck et al., 2019). In the frontal cortex, a hypomethylation was observed. No genome-wide studies have been conducted on nicotine yet, which may be surprising considering the large population of women exposed to that substance during pregnancy.

## 8. Discussion and perspectives

Following the kinetics that unfold during development, somatic cells display a heavily methylated landscape largely shared across cell types (Bird, 2002; Smith et al., 2024). While neurons are post-mitotic cells that have long been considered to exhibit static DNAm patterns, they nevertheless express Dnmt and Tet enzymes at relatively high levels (Lister et al., 2013; Szwagierczak et al., 2010; Li et al., 2014). As illustrated in the present review, multiple pieces of evidence now strongly argue for the existence of a significant degree of DNAm plasticity in these cells in relation to SUD (Lister and Mukamel, 2015).

Collectively, available results suggest that this plasticity may be different across drugs of abuse or brain regions. However, direct genome-wide comparisons specifically addressing these differences remain to be conducted, and represent an immediate perspective. Such efforts are expected to generate new insight by differentiating drug-specific mechanisms (which reflect their distinct pharmacological targets) from those that may be common to multiple drugs, and therefore at the core of the behavioral dysregulation defining SUD.

Early genome-wide studies in the neuroscience field used methodological approaches that focused on promoter regions. Inspired by studies conducted in the cancer field, this reflected the historical assumption that adaptations at these regions were primarily responsible for changes in transcriptional activity. In contrast, the present systematic review clearly indicates that when newer unbiased methods were used, most of the observed DNAm changes occurred outside these regions (Fig.7e), both within and far away from genes. Our own recent work on opioids further reinforces this notion (Falconnier et al.,2025). Therefore, future studies in the field should shift away from a promoter-centric view of gene regulation in the rodent brain.

As already mentioned, this notion is consistent with the fact that the magnitude of gene expression changes observed in rodent models of SUD is typically low compared to those observed in relation to developmental or pathological changes in cell identity. This may in part reflect the biological nature of brain tissue, which is mostly composed of post-mitotic cells expressing a relatively constrained transcriptomic profile. However, it also likely reflects the technical limitation of bulk tissue analyses. Accordingly, the brain consists of a wide variety of cell types, and deciphering the cell-type specific DNAm changes and their functional effects represent an important challenge for the coming years.

Another important consideration is that the interplay of drug-induced DNAm changes with other regulatory factors, such as histone marks or transcription factors, is largely unresolved. Furthermore, how these factors interact to drive changes in transcriptional networks remains unclear, as correlations among DNAm and the transcriptome were repeatedly subtle across the studies reviewed in the present work. Overall, this calls for the generation of multiomic and genome-wide datasets in the field. Ideally, these data should be analyzed using computational methods that go beyond the mere aggregation of sparse differences observed for individual omic layers, and leverage co-expression networks, dimensionality reduction and nonlinear methods of data integration (Mokhtari et al. 2025; Mokhtari et al. 2022).

Finally, new tools and knowledge about DNAm enzymes should drive future work. To date, the localization of these enzymes remains difficult to measure with good genomic resolution, and available data for e.g. Dnmt3a suggests diffuse patterns with fluctuations of little amplitude (Stroud et al., 2020, 2017; Gu et al., 2022). New tools such as knockin mouse lines with tagged enzymes (Gu et al., 2022) should help researchers precisely map the genomic localisation of Dnmt and Tet, and identify possible changes in pathological contexts. Also, recent evidence suggests that Dnmt1 has significant *de novo* methylation activity (Haggerty et al., 2021; Li et al., 2018), which may have implications in brain tissue, where this enzyme remains readily expressed by post-mitotic neurons, although at lower levels than Dnmt3a (Lister et al., 2013). Another aspect is that 2 domains have recently been characterized in the C-terminal region of Dnmt3a, in addition to the 2 classical catalytic and ATRX–DNMT3–DNMT3L (ADD) domains responsible for methylating DNA and interactions with H3K4 methylation, respectively (Ooi et al., 2007). The first, PWWP, is involved in the dual recognition of H3K36me2 and H3K36me3, and in shaping intergenic patterns of DNAm in mouse mesenchymal or embryonic stem cells (Weinberg et al., 2019). Interestingly, this interaction accounts in the brain for the postnatal emergence of 1 Mb-sized domains of 5mCH that follow the spatial organization of chromatin (into topologically associating domains, TAD; (Hamagami et al., 2023)), and represent functional units for transcriptional regulation (Clemens et al., 2020). The second domain corresponds to a ubiquitin-dependent region, present in the Dnmt3a1 but not the Dnmt3a2 isoform, which have different roles in learning and memory (Oliveira et al., 2016; Oliveira et al., 2012). This domain was recently found to mediate recruitment of Dnmt3a at nucleosomes marked by H2AK119ub (Weinberg et al., 2021; Gu et al., 2022). Surprisingly, these 2 domains, the H3K36me2/3 and H2AK119ub histone marks, as well as large mCH domains have not been investigated in relation to SUD phenotypes. Determining whether and how they may contribute to pathophysiology represent important avenues for future work.

