## Supplementary material for "The DNA methylation enzymatic machinery in substance use disorders: a systematic review"

Pierre-Eric Lutz, INCI UPR 3212, 8 allée du général Rouvillois, 67000 Strasbourg, France

**Legends of Supplementary Figures**

**Supplementary Figure 1. Global measures of changes in DNA methylation levels (averaged throughout the genome) following exposure to drugs of abuse.** Rows correspond to brain regions and columns to drugs of abuse. Colors refer to changes in global DNA methylation levels: yellow indicates that an increase was observed, blue a decrease, and grey that there were no detectable changes. The asterisk indicates that a measure of 5hmC level was performed. Abbreviations: Amy, amygdala; CPu, caudate putamen; Hip, hippocampus; NAc, nucleus accumbens; OFC, orbitofrontal cortex; PFC, prefrontal cortex; Str, striatum; VTA, ventral tegmental area.

**Supplementary Figure 2. Candidate gene studies on cocaine.** The table summarizes the brain region and genes that were investigated. Yellow and blue indicate a decrease or an increase in DNA methylation following cocaine exposure, respectively, while grey indicates no significant changes. Upward or downward arrows indicate an increase or a decrease in gene expression, respectively, while "-" indicates no significant changes. Abbreviations: Cc, corpus callosum; CPu, caudate putamen; Hip, hippocampus; LHb, lateral habenula; NAc, nucleus accumbens; PFC, prefrontal cortex; dmPFC, dorsomedial prefrontal cortex; Str, striatum.

**Supplementary Figure 3. Candidate gene studies on opioids.** The table summarizes the brain region and genes that were investigated. Yellow and blue indicate a decrease or an increase in DNA methylation following cocaine exposure, respectively, while grey indicates no significant changes. Upward or downward arrows indicate an increase or a decrease in gene expression, respectively, while "-" indicates no significant changes. Abbreviations: Cb, cerebellum; Ce, cerebral cortex; Hip, hippocampus; Hyp, hypothalamus; IC, inferior colliculus; Mi, midbrain; MO, medulla oblongata; NAc, nucleus accumbens; mPFC, medial prefrontal cortex; Po, pons; SC, spinal cord; Th, thalamus; VTA, ventral tegmental area.

**Supplementary Figure 4. Candidate gene studies on methamphetamine.** The table summarizes the brain region and genes that were investigated. Yellow and blue indicate a decrease or an increase in DNA methylation following cocaine exposure, respectively, while grey indicates no significant changes. Upward or downward arrows indicate an increase or a decrease in gene expression, respectively, while "-" indicates no significant changes. Abbreviations: DG, dentate gyrus; Hip, hippocampus; NAc, nucleus accumbens; PFC, prefrontal cortex; Str, striatum.
