## Supplementary Figure 1 for "The DNA methylation enzymatic machinery in substance use disorders: a systematic review"

**Figure 4**

| | Cocaine | Alcohol | Methamphetamine | Opioids | Nicotine |
|---|---|---|---|---|---|
| Whole brain | Fragou, 2013<br>Chao, 2014 * | | | Fragou, 2013<br>Chao, 2014 * | Nguyen, 2019 |
| NAc | Tian, 2012<br>Feng, 2015 *<br>Wright, 2015<br>Anier, 2018 * | Niinep, 2021 *<br>Barbier, 2015 | Mychasiuk, 2013 | Tian, 2012 | Mychasiuk, 2013 |
| PFC | Tian, 2012<br>Saad, 2021 | Yang, 2021<br>Barbier, 2015 | Mychasiuk, 2013<br>Gonzalez, 2018 | Tian, 2012 | Mychasiuk, 2013<br>Buck, 2019 |
| OFC | | | | | Mychasiuk, 2013 |
| Str | | | | Joanna, 2017 | Buck, 2019 |
| VTA | | | | Fan, 2019 * | |
| Hip | | Yang, 2021<br>Barbier, 2015 | | Fan, 2019 *<br>Fan, 2021<br>Chen, 2019<br>Fan, 2021 | Nguyen, 2019 |
| CPu | Saad, 2021 | | | | |
| Amy | | Yang, 2021<br>Barbier, 2015 | | | |
| Neocortex | | Bottom, 2020 | | | |

| **Legend** |
|---|
| Hypomethylation |
| Hypermethylation |
| No changes |
| * 5hmc assayed |
