## Supplementary Figure 2 for "The DNA methylation enzymatic machinery in substance use disorders: a systematic review"

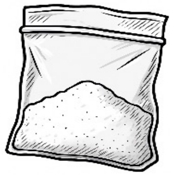

### Cocaine

| | FosB | Crem | Pp1c | | Cdkl5 | Auts2 | Caln1 | miR-124 | GLT-1 | Mbp | Plp1 | Sox10 | Homer2 | orx-R | Bdnf | A2AR | Taar7b |
|---|---|---|---|---|---|---|---|---|---|---|---|---|---|---|---|---|---|
| | | | Pp1cβ | Ppp1cc | | | | | | | | | | | | | |
| Ajonijebu,2017 | PFC | | | | | | | | | | | | | | | | |
| | Hip | | | | | | | | | | | | | | | | |
| Anier,2010 | | | NAc ↓ | | | | | | | | | | | | | | |
| Pol Bodetto, 2013 | | | PFC | | | | | | | | | | | | | | |
| | | | CPu ↑ | | | | | | | | | | | | | | |
| Vaher, 2020 | | | | NAc ↓ | | | | | | | | | | | | NAc ↓ | NAc ↑ |
| | | | | LHb ↓ | | | | | | | | | | | | LHb ↓ | LHb ↑ |
| Carouge, 2010 | | | | | Str ↓ | | | | | | | | | | | | |
| Engmann, 2017 | | | | | | NAc ↑ | | | | | | | | | | | |
| Guo, 2016 | | | | | | | | Whole brain ↓ | | | | | | | | | |
| Kim, 2018 | | | | | | | | | NAc ↓ | | | | | | | | |
| Nielsen, 2012 | | | | | | | | | | Cc | Cc | Cc | | | | | |
| Ploense, 2018 | | | | | | | | | | | | | dmPFC ↑ | | | | |
| Saad,2019 | | | | | | | | | | | | | | PFC ↑ | | | |
| Tian, 2016 | | | | | | | | | | | | | | | NAc ↑ | | |

| Methylation | Gene expression |
|---|---|
| Hypomethylation | ↑ : Upregulation |
| Hypermethylation | ↓ : Downregulation |
| No changes | - : No changes |
