## Supplementary Figure 3 for "The DNA methylation enzymatic machinery in substance use disorders: a systematic review"

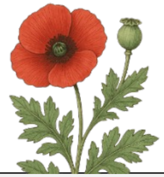

### Opioids

| | Drd2 | Erg1 | Erg2 | Oprm1 | Gr17 | Crhbp | Bdnf | Comt | Il1b | Il6 | Nr3c1 | Tnf | L1-orf | L1-5utr | Camkk1 |
|---|---|---|---|---|---|---|---|---|---|---|---|---|---|---|---|
| Rodrigues, 2011 | NAc ↓ | | | | | | | | | | | | | | |
| Imperio, 2018 | | mPFC ↓ | mPFC ↓ | mPFC - | | | | | | | | | | | |
| | | NAc ↓ | NAc ↓ | NAc - | | | | | | | | | | | |
| Zhu, 2017 | | | | | Hip ↑ | | | | | | | | | | |
| MacFalls, 2016 | | | | | | VTA ↑ | | | | | | | | | |
| Barrow, 2017 | | | | | | | Po | Po (chronic) | Po | Po (acute) | Po | Po | Po | Po | |
| | | | | | | | Hip | Hip | Hip (acute) | Hip | Hip (chronic) | Hip | Hip | Hip | |
| | | | | | | | Cb | Cb | Cb | Cb | Cb (acute) | Cb | Cb | Cb | |
| | | | | | | | MO | MO | MO (acute) | MO | MO | MO | MO | MO | |
| | | | | | | | Ce | Ce | Ce | Ce | Ce | Ce | Ce | Ce | |
| | | | | | | | Hyp | Hyp | Hyp | Hyp | Hyp | Hyp | Hyp | Hyp (chronic) | |
| | | | | | | | Mi | Mi | Mi | Mi | Mi | Mi | Mi | Mi | |
| | | | | | | | IC | IC | IC | IC | IC | IC | IC | IC | |
| | | | | | | | SC | SC | SC | SC | SC | SC | SC | SC | |
| | | | | | | | Th | Th | Th | Th | Th | Th | Th | Th | |
| Chen, 2019 | | | | | | | | | | | | | | | Hip ↑ |

| **Methylation** | **Gene expression** |
|---|---|
| Hypomethylation | ↑ : Upregulation |
| Hypermethylation | ↓ : Downregulation |
| No changes | - : No changes |
