## Supplementary Figure 4 for "The DNA methylation enzymatic machinery in substance use disorders: a systematic review"

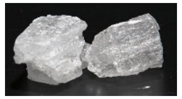

### Methamphetamine

| | α-syn | Arc | Erg2 | Fos | Nr4a1 | Klf10 | Syn | Bdnf IV | Glua1 | Glua2 | Kcna1 | Kcna3 | Kcnn1 | Line-1 | Pabp1 |
|---|---|---|---|---|---|---|---|---|---|---|---|---|---|---|---|
| Biagioni, 2019 | Str ↑ | | | | | | | | | | | | | | |
| Cheng, 2015 | | PFC ↓<br>Hip ↓ | PFC ↓<br>Hip ↓ | PFC ↓<br>Hip ↓ | PFC ↓<br>Hip ↓ | PFC -<br>Hip - | | | | | | | | | |
| Fan, 2020 | | | | | | | Hip ↑<br>PFC ↓ | | | | | | | | |
| Iamjan, 2021 | | | | | | | | Hip ↑<br>PFC - | | | | | | Hip, Str, PFC | |
| Jayanthi, 2014 | | | | | | | | | Str ↓ | Str ↓ | | | | | |
| Jayanthi, 2019 | | | | | | | | | | | NAc ↑ | NAc ↑ | NAc ↑ | | |
| Moszczynska, 2015 | | | | | | | | | | | | | | Str ↑<br>DG ↑ | |
| Moszczynska, 2016 | | | | | | | | | | | | | | | DG ↑ |
| Salehzadeh, 2020 | | | | | | | | PFC ↑ | | | | | | | |

| Methylation | Gene expression |
|---|---|
| Hypomethylation | ↑ : Upregulation |
| Hypermethylation | ↓ : Downregulation |
| No changes | - : No changes |
